# New insights into the echinocandin resistance in *Candida* spp. in the clinical setting

**DOI:** 10.1101/2024.04.29.591739

**Authors:** Antoine Gedeon, Yasmine Kalboussi, Jeanne Bigot, Vicky Le, Eric Dannaoui, Sandra Vellaissamy, Marie Antignac, Stéphanie Petrella, Christophe Hennequin, Juliette Guitard

**Affiliations:** Université Paris Cité, Institut Pasteur, Unité de Microbiologie Structurale, CNRS UMR3525, F-75015 Paris, France; Sorbonne Université, AP-HP, Hôpital Saint-Antoine, Service de Parasitologie-Mycologie, F-75012 Paris, France; Sorbonne Université, Inserm, Centre de Recherche Saint-Antoine, CRSA, AP-HP, Hôpital Saint-Antoine, Service de Parasitologie-Mycologie, F-75012 Paris, France; Unité de Parasitologie-Mycologie, Hôpital Necker - Hôpital Européen Georges Pompidou, AP-HP, Faculté de Médecine, Université de Paris, F-75015 Paris, France; Sorbonne Université, AP-HP, Hôpital Saint-Antoine, Service de Pharmacie, F-75012 Paris, France

**Keywords:** *Candida*, echinocandin resistance, (1,3)-β-D-Glucan synthase, fitness, molecular modelling

## Abstract

Despite a huge consumption of echinocandins, the emergence of resistance in *Candida spp* has remained overall limited. Here, we depicted the epidemiology of *Candida* spp in our center face to the echinocandins’ consumption. We postulate new hypotheses that may explain the shaping of candines resistance in the clinical setting. Epidemiology of *Candida* infections and echinocandin consumption were evaluated in our center over 12 years (2006-2018). Glucan synthase genes (*fks1, fks2)* were sequenced. The *in vitro* fitness was assessed for couple of isogenic strains, of which one was resistant. Finally, the modelling of *Candida* FKS proteins was realized. Despite a three-fold increase in echinocandin consumption, no significant emergence of resistance was observed. In *Candida albicans, fks1* mutations affect the three-dimensional conformation of the glucan synthase, which may result in altered export of the glucan chain, which could explain the observed reduced fitness. In contrast, cross-complementation between FKS1 and FKS2 in *Nakaseomyces glabratus* might circumvent the negative impact on fitness.

**Conclusions:** Our results support the fact that *N*. glabratus is more prone to the spread of echinocandin-resistant strains. The link between mutations in the glucan synthase and fitness of *Candida* strains might explain the difference in the species-specific emergence of echinocandins-resistant strains.

## Introduction

Invasive candidiasis is a life-threatening infection that needs early and appropriate antifungal therapy. Guidelines from the Infectious Diseases Society of America and the European Society of Clinical Microbiology and Infectious Diseases recommend an echinocandin as first-line therapy for invasive candidiasis [1,2]. Due to their safety profile and fungicidal activity, those drugs have been extensively used for therapeutic, empiric or pre-emptive use. Concomitantly, the number of reports either on *Candida* strains with acquired reduced-echinocandin-susceptibility (RES), with the noticeable relative predominance of *Nakaseomyces glabratus* (formerly *Candida glabrata)* [3,4] or the change in the epidemiology in favour of *Candida* species naturally less susceptible to echinocandins, notably *Candida parapsilosis* increased gradually [5–7].

From a mechanistic point of view, it has been speculated that echinocandins intercalate in the phospholipid bilayer of the plasma membrane and inhibit the (1,3)-β-D-Glucan synthase complex (GSC). The GSC consists of a catalytic unit (FKS1, FKS2 and FKS3 proteins) responsible for the (1,3)-β-D-Glucan chain elongation and a regulatory unit (RhoGTPase) [8]. FKS1 seems to be the primary source of glucan polymers in *Candida albicans*, since the knock-out of *fks1* is lethal in this species [9]. In contrast, in *N. glabratus* and *Saccharomyces cerevisiae*, FKS1 and FKS2 appear to be redundant for the synthesis of the (1,3)-β-D-Glucan [10,11]. Despite increasing knowledge, the precise molecular organization of the GSC remains incompletely elucidated for the different *Candida* species.

Reduced-echinocandin-susceptibility, whether it is natural or acquired, mostly relies on nucleotide mutations within two hot-spot (HS) regions in the *fks1* or *fks2* gene [5,12]. Up to now, few studies have investigated the impact of acquired *fks* mutations on the virulence of clinical strains of *Candida* [13]. Using both laboratory-derived and clinical *C. albicans*-resistant strains, *fks1* mutations have been associated with an impaired growth rate and decreased virulence in animal models of invasive candidiasis [14]. In contrast, a conserved fitness has been documented in various animal models for echinocandin-resistant clinical *N. glabratus* strains harbouring mutations in *fks2* HS regions [15,16].

In this retrospective study, we reviewed the epidemiology of *Candida* species in our center face to the echinocandins consumption over 12 years (2006-2018). In the second part, isolates with acquired RES were characterized and the fitness of pairs of isogenic isolates (echinocandin-susceptible and resistant) was assessed comparatively. Finally, molecular modelling of FKS1 and FKS2 allowed us to hypothesize on the impact of mutations associated with RES on the fitness of the strains in a species-specific manner.

## Results

### Trends in Candida epidemiology and echinocandins consumption

Between 2006 and 2013, the echinocandins’ consumption increased three-fold, then remained overall stable in subsequent years (Figure 1A). Concomitantly, an increase in *C. parapsilosis* frequency was noted from 1.5% in 2006 to 8.3% in 2013, and further to 10.9% in 2018 (*p*=0.01, Pearson coefficient 0.66 [CI95 0.17-0.89]) (Figure 1B). The prevalence of patients with a RES strain varied non-significantly between 0% and 4% each year (Figure 1C). In total, among 2,754 strains identified and for which the echinocandin susceptibility was determined among 2268 patients, we detected 21 (0.8%) RES strains collected from 18 (0.8%) patients, suffering from various forms of candidiasis.

**Figure 1:**
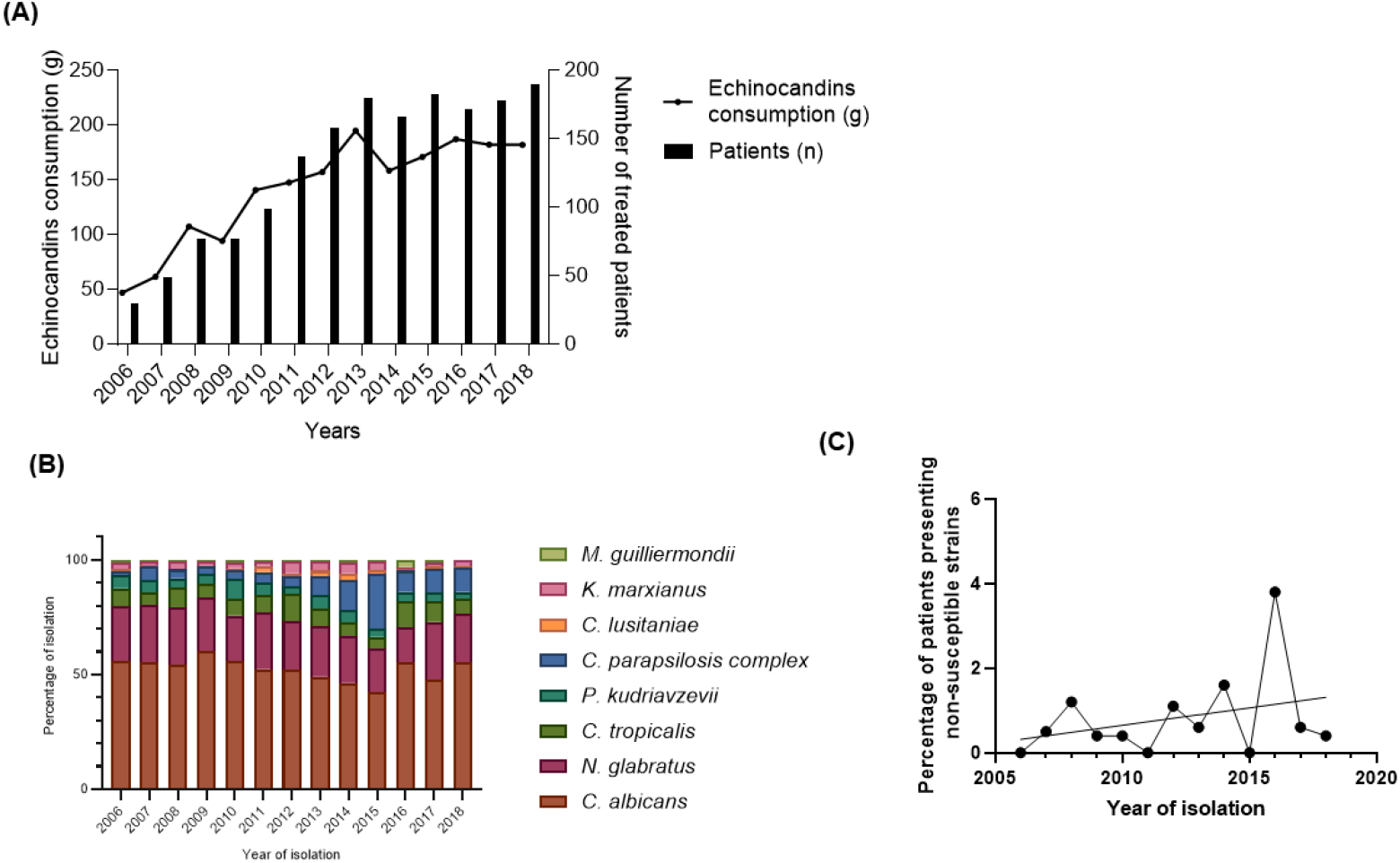
Epidemiology and echinocandins consumption in our University Hospital from 2006 to 2018. (A), Echinocandin consumption in g/year and number of patients treated by echinocandin/year. (B), Epidemiological evolution of *Candida* species distribution. Non-significant evolution of *C. albicans* (*r*= -0.5, *p*=0.09), *N. glabratus* (*r*=-0.5, *p*=0.08), C. *tropicalis* (*r*=0.2, *p*=0.6), *P. kudriavzevii* (*r*=-0.5, *p*=0.1), *C. lusitaniae* (*r*=0.2; *p*=0.5), *K. marxianus*(*r*=-0.07, *p*=0,.8), *M. guilliermondii* (*r*=0.3, *p*=0.3), and significant increase in *C. parapsilosis* complex percentage of isolation (*r*=0.7, *p*=0.01) as analyzed with a Pearson multiple correlation test. (C), Percentage of patients presenting non-susceptible strains according to the year.

### Patients with acquired RES strains

Clinical and biological charts were available for 14 of the 18 patients infected with an acquired RES strain (Table 1). Species involved in these cases were *N. glabratus* (*n*=6), *C. albicans* (*n*=3) and *Kluyveromyces marxianus* (formerly *Candida kefyr*) (*n*=3). For 17 of the 18 strains, we found non-synonymous mutations in HS1 or HS2 of *fks* genes (Table 1).

**Table 1:**
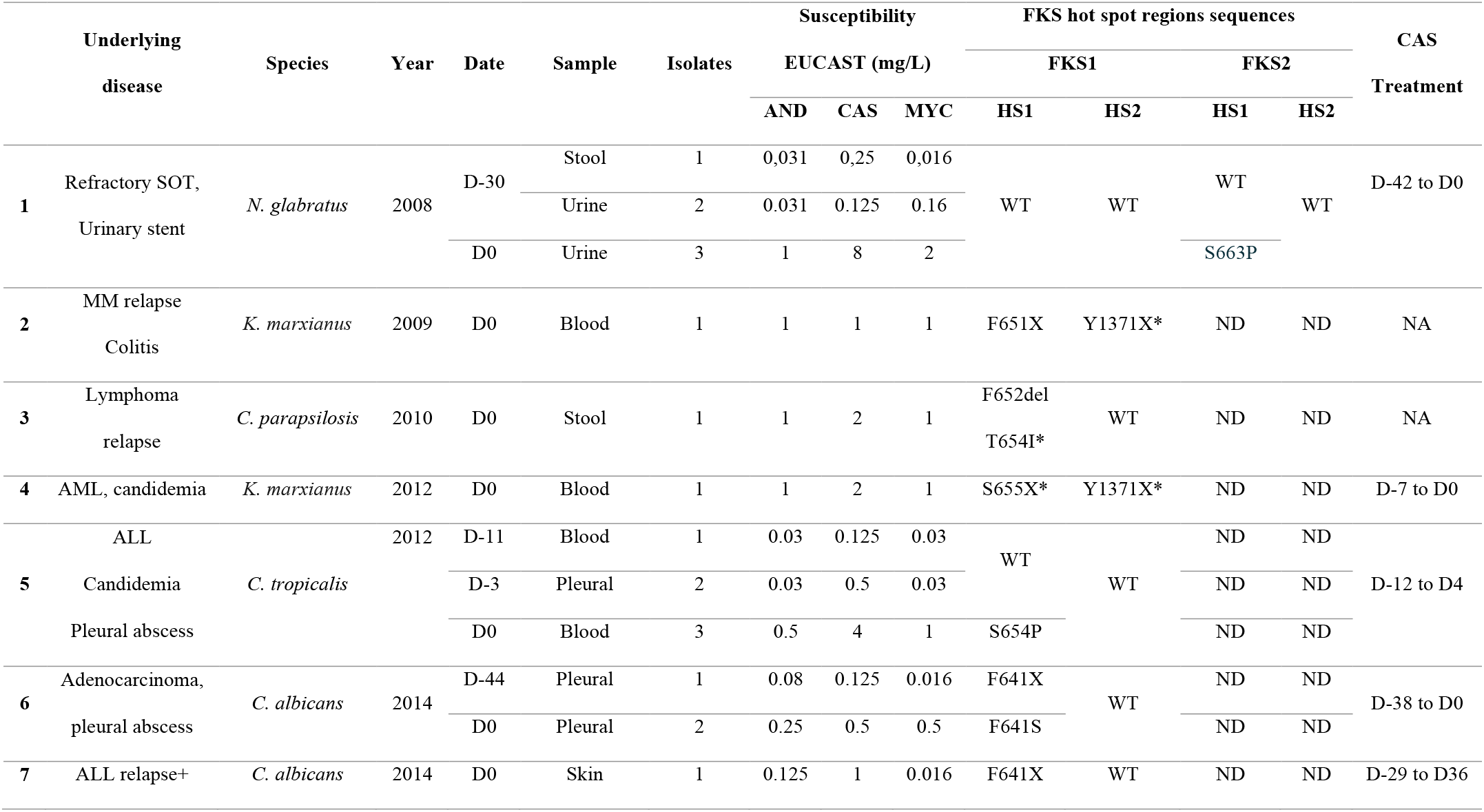
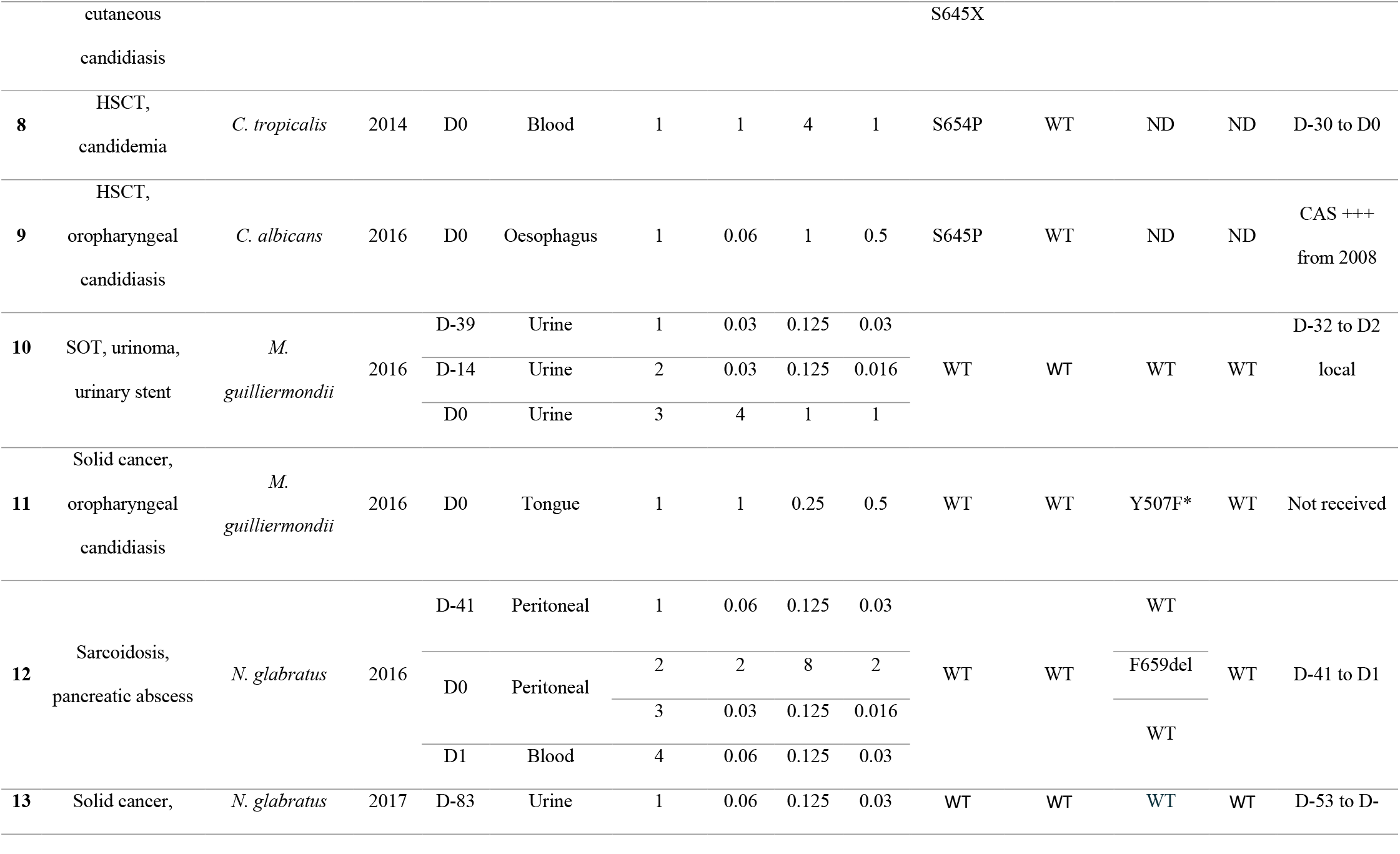
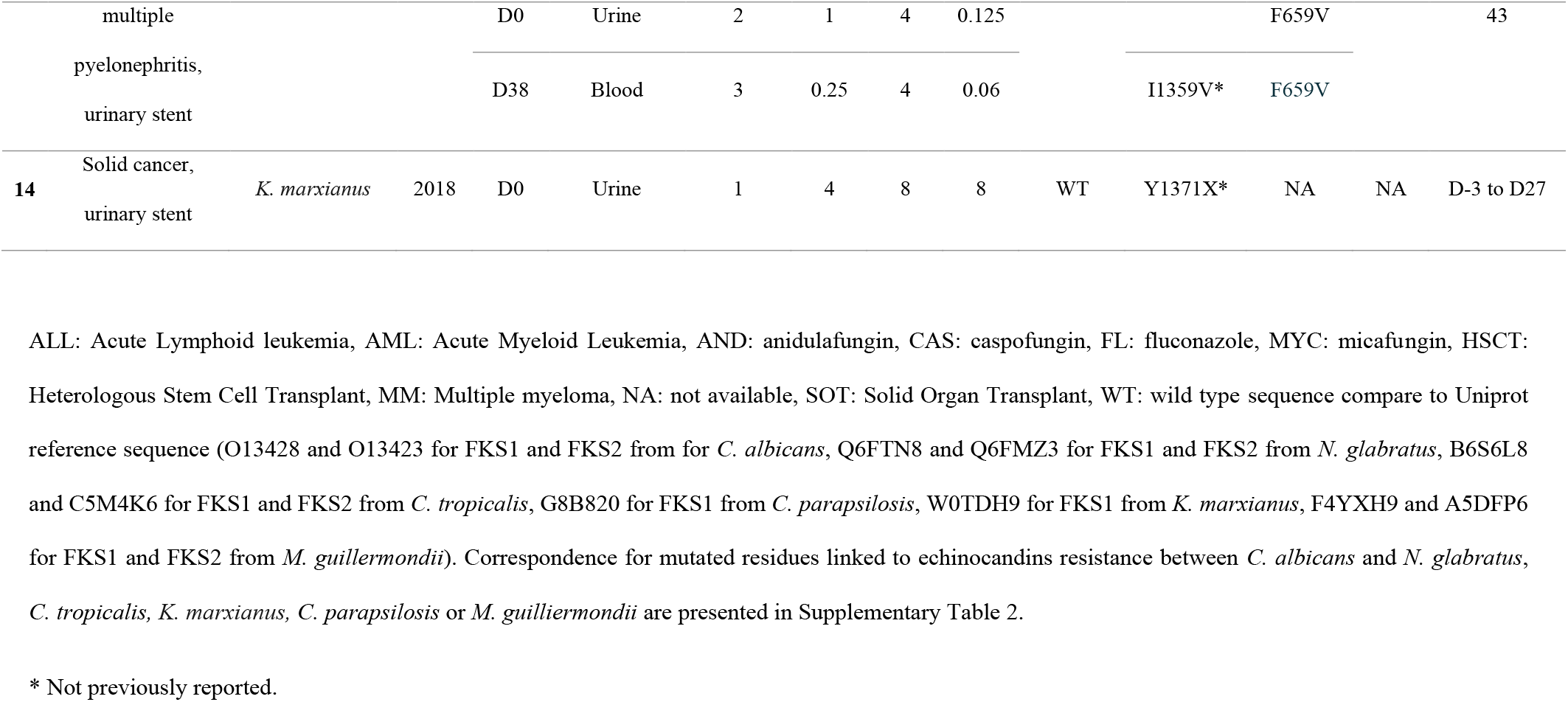
Characteristics of patients and strains of patients infected by resistant strains.

Among 12 patients with available information, one had previously received multiple echinocandin courses while echinocandin therapy was ongoing for 10 others. In these cases, the RES strain was isolated after a median of 29.5 days following the initiation of the echinocandin-based treatment (*n*=10; range: 3-41 days) (Table 1). Interestingly, among the 14 patients with available data, 11 (78.6%) had biofilm-associated infections characterized by a heavy fungal burden (Table 1).

### RES-Candida isolate fitness

Five echinocandin-susceptible and six RES isolates with *fks* mutations collected from 5 patients were available for testing. Microsatellite typing confirmed the isogenic character of those couples of isolates, supporting the hypothesis of acquisition of resistance rather than superinfection with a different strain (data not shown). We found two different fitness patterns for susceptible *versus* RES isolates (Table 2). The first pattern (“type 1”) showed RES isolates with a reduced fitness compared to their isogenic susceptible counterparts. This pattern was observed for *C. albicans* and *C. tropicalis* strains, all harbouring mutations in the HS1 region of *fks1* (Table 2). The second pattern (“type 2”) showed a similar fitness for the susceptible and the resistant isolates in YPD medium, but an increased fitness for resistant isolates in stress conditions. This pattern was observed in three *N. glabratus* strains harbouring mutations in the HS1 region of *fks2* (Table 2).

**Table 2:**
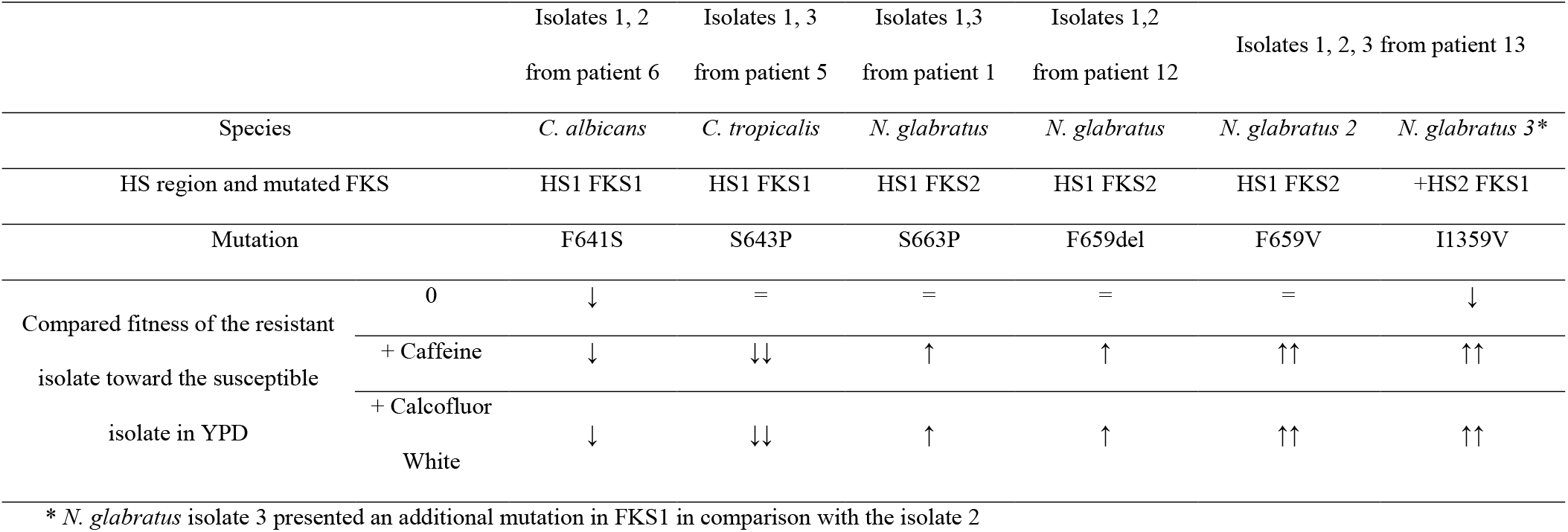
*In vitro* characterization of *Candida* fitness of the resistant isolate compared to the isogenic susceptible isolate for each patient with isogenic isolates available. Species are indicated, as well as the FKS mutation observed in the resistant isolate. Comparisons of fitness of the resistant isolate toward the susceptible isolate in YPD as well as in stress media: YPD + caffeine and YPD + Calcofluor White are presented.

### Mapping of mutated residues associated with echinocandin resistance in FKS1 and FKS2

Three orthologues of FKS proteins have been described, with FKS1 and FKS2 being responsible for (1,3)-β-D-Glucan synthesis in fungi. FKS1 is highly conserved among the *Saccharomycetales* including genetically distant species such as *C. albicans* and *N. glabratus* (similarity >73%). Noteworthy, FKS2 from *N. glabratus* also exhibits a similarity rate of 72% with *C. albicans* FKS1 while the percentage of similarity was lower at 53% for *C. albicans* FKS2 (Supplementary Figure 1).

To investigate the role of mutations in FKS1 or FKS2 associated with echinocandin resistance on the protein’s structure and a possible link with their function, we modelled FKS1 and FKS2 monomers and mapped the mutated residues (Figure 2A, B). The modelled structures showed high-level confidence and low residue position errors (Supplementary Figure 2). As expected, FKS1 and FKS2 comprise of two distinct domains. The first is a series of 17 contiguous α-helixes which constitutes the transmembrane domain; the second is a globular domain corresponding to the catalytic intracytoplasmic domain where glucan synthesis occurs. Importantly, all the mapped mutations in FKS1 (except M1235) and FKS2 are localized within the transmembrane domains.

**Figure 2:**
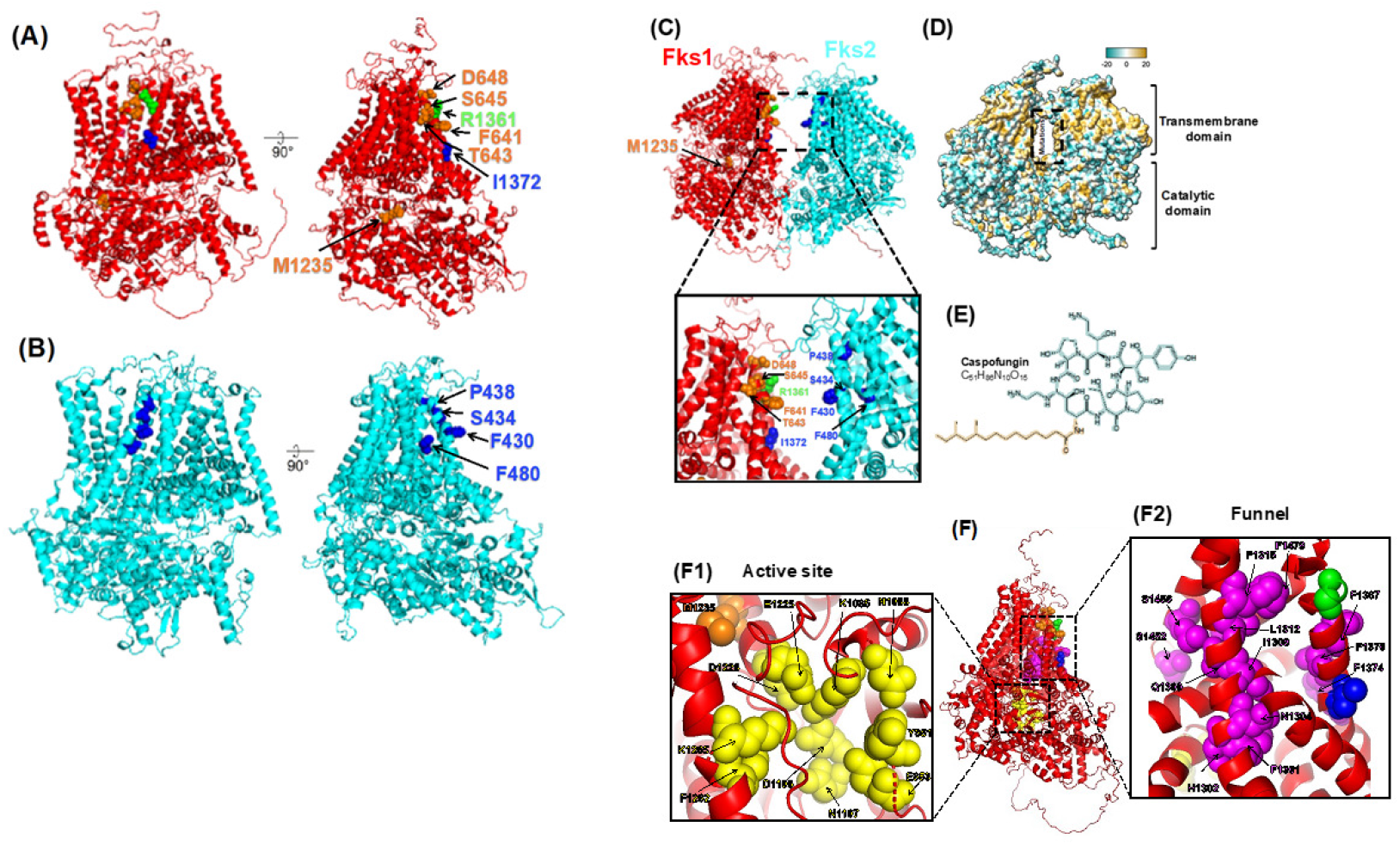
Structural models of *C. albicans* FKS1, FKS2 and FKS1:FKS2 generated with AlphaFold2. (A-C) Mapping of mutations on models of *C. albicans* FKS1 monomer (A), FKS2 monomer (B), and FKS1:FKS2 dimer (C). Residues were numbered according to *C. albicans* sequences. FKS1 and FKS2 monomers are colored red and cyan respectively. Mutated residues linked to echinocandin resistance are highlighted in spheres and colored as follows: in orange for residues identified in *C. tropicalis* (F641, T643, S645; D648 and M1235), in green for residues identified in *C. albicans* (R1361, F461 and S645; note that F461 and S645 were also described in *C. tropicalis* mutants, therefore colored in orange for simplicity), in blue for residues identified in *N. glabratus* (F430, S434, P438, F480, I1372). (D, E) Hydrophobicity map of FKS1:FKS2 surface (D) and caspofungin (E). Hydrophilic (lipophobic, polar) moieties are colored in cyan, and hydrophobic (lipophilic, nonpolar) moieties are colored in yellow gold. Residue numbering correspondence between the three species is summarized in supplementary table 2. (F) AlphaFold2 model of *C. albicans* FKS1 monomer showing residues of the active site (zoom in F1) and residues of the product transfer funnel linking the active site to the extra-cytoplasmic compartment (zoom in F2). Residues were numbered according to *C. albicans* sequences. Mutated residues linked to echinocandin resistance are highlighted in spheres and colored as described in Figure 2. Residues of the active site or implicated in product export funnel are highlighted in spheres and colored yellow and magenta respectively. Residue numbering correspondence between *S. cerevisiae* and *C. albicans* is summarized in supplementary table 3.

To evaluate a possible interaction between FKS1 and FKS2, we then modelled an FKS1:FKS2 dimer and mapped all the described mutations within the binary complex. This showed a relevant fit between FKS1 and FKS2 and a striking localization of mutations at the FKS1:FKS2 interface within adjacent transmembrane domains of each monomer (Figure 2C). This interface is strongly hydrophobic (Figure 2D) and could interact with the hydrophobic echinocandin fraction (Figure 2E).

Using recent structural investigations on FKS1 [17,18], we additionally showed that FKS mutations are not implicated in the substrate binding or catalysis (zoom F1 in Figure 2F) but rather hamper the exit of the funnel to export (1,3)-β-D-Glucan products into the extra-cytoplasmic compartments (Zoom F2 in Figure 2F).

## Discussion

During the period 2006-2018, a huge increase in echinocandins consumption was observed in our center. We faced a moderate increase in *C. parapsilosis* and *M. guilliermondii* frequency, two species known to exhibit naturally high echinocandins MIC [19]. In contrast, the emergence of acquired RES remains very limited (<4% of patients each year). Our results are in line with a national French survey, conducted from 2002 to 2009, showing an increase of *C. parapsilosis* frequency in candidemia from 13 to 31% in parallel to the increase of echinocandins’ consumption, whereas there was no emergence of acquired-RES for other main pathogenic species [6].

In our study, most of the patients infected with or harbouring an acquired RES strain had a biofilm-associated infection characterized by a heavy fungal burden and a previous echinocandin treatment lasting for a median of 29.5 days. These findings are consistent with data from a surveillance program which found 30 days of echinocandin exposure preceding the emergence of acquired RES strains [20]. A heavy fungal burden may have favoured the emergence of mutants under antifungal pressure as mutations in *fks* genes were detected in almost all the RES isolates.

To test the hypothesis that this low and stable rate of acquired-RES strains could be due to the reduced fitness of the *fks* mutant isolates, we compared couples of isogenic isolates. Two profiles were observed according to the *Candida* species. RES-*C. albicans* isolates with *fks1* mutations presented a reduced fitness compared to the isogenic susceptible isolate. A reduced virulence for *C. albicans* strains harbouring mutations in the HS1 region of FKS1 has also been observed in *Toll*-Deficient *Drosophila melanogaster* and murine models [14]. These data confirm that FKS1 mutations can impact the fitness of *C. albicans*, adding evidence to a previous study suggesting that FKS2 is not efficient for the synthesis of (1,3)-β-D-Glucan in this species [9]. In contrast, RES-*N. glabratus* isolates with *fks2* mutations presented a fitness similar, or even increased, compared to the isogenic susceptible isolate. Similarly, two studies using murine and *Galleria mellonella* models showed that RES-*N. glabratus* strains harbouring mutations in the HS1 region of FKS2 did not exhibit any decrease in their fitness or virulence [15,16]. This crucial difference between species might explain the relative predominance of *N. glabratus* in papers reporting on echinocandin-acquired resistance, in comparison to other pathogenic species, including *C. albicans*.

A possible explanation for this difference led us to hypothesize a pivotal role of FKS2 in maintaining a conserved glucan synthesis in *N. glabratus* mutants. We showed that FKS1 from all *Saccharomycetales* and FKS2 from *N. glabratus* shared a high percentage of similarity, whereas FKS2 from other species, including *C. albicans*, is more divergent. This supports a functional redundancy of FKS1 and FKS2 in *N. glabratus* [10], a phenomenon that doesn’t exist in other *Candida* species evaluated in this study, for which FKS1 seems essential [9].

Despite being essential for maintaining *ad hoc* concentrations of (1,3)-β-D-Glucan in the cell wall and being a therapeutic target of antifungal drugs, very little is known about the three-dimensional structure of GSC in the *Saccharomycetales*. Notably, no specific binding site(s) have yet been identified for echinocandins. It has been proposed that echinocandins intercalate in the phospholipid bilayer of the plasma membrane *via* their lipid tail and inhibit the GSC [5]. Two recent independent functional analysis and cryo-electron microscopy studies in *S. cerevisiae* suggested that, after purification conditions maintaining the membrane architecture, FKS1 exists in a monomeric form [17,18]. Modelling FKS1 and FKS2 monomers shows that all mutations are present in the previously described HS regions which have been speculated to be implicated in interactions with echinocandins [5]. Otherwise, cryo-electron tomography on a *N. glabratus* strain that overexpressed FKS1, suggested an organization of GSC as a high-order hexamer formed by FKS1 and/or FKS2 proteins [21]. In our study, when modelling included an FKS1:FKS2 dimer, we observed a hydrophobic pocket at the FKS1/FKS2 interface which could serve as both a binding pocket for the hydrophobic moiety of echinocandins and an exit funnel for the glucan chain as suggested by Hu et al [17]. Interestingly, all the described mutations in FKS proteins concerned residues involved in the constitution of this pocket, making our hypothesis relevant (Figure 3). Our results and those in the literature, lead us to hypothesize that mutations in *C. albicans* FKS1, the unique protein responsible for glucan synthesis in this species, both affect the binding of echinocandins, leading to RES, but also the exit of the glucan chain, resulting in a reduced fitness of mutant strains. In contrast, in *N. glabratus*, where FKS1 and FKS2 may exist as polymers, mutations, mostly seen in FKS2, impact the echinocandins binding but not the exit of the glucan chain therefore maintaining the strains’ fitness.

**Figure 3:**
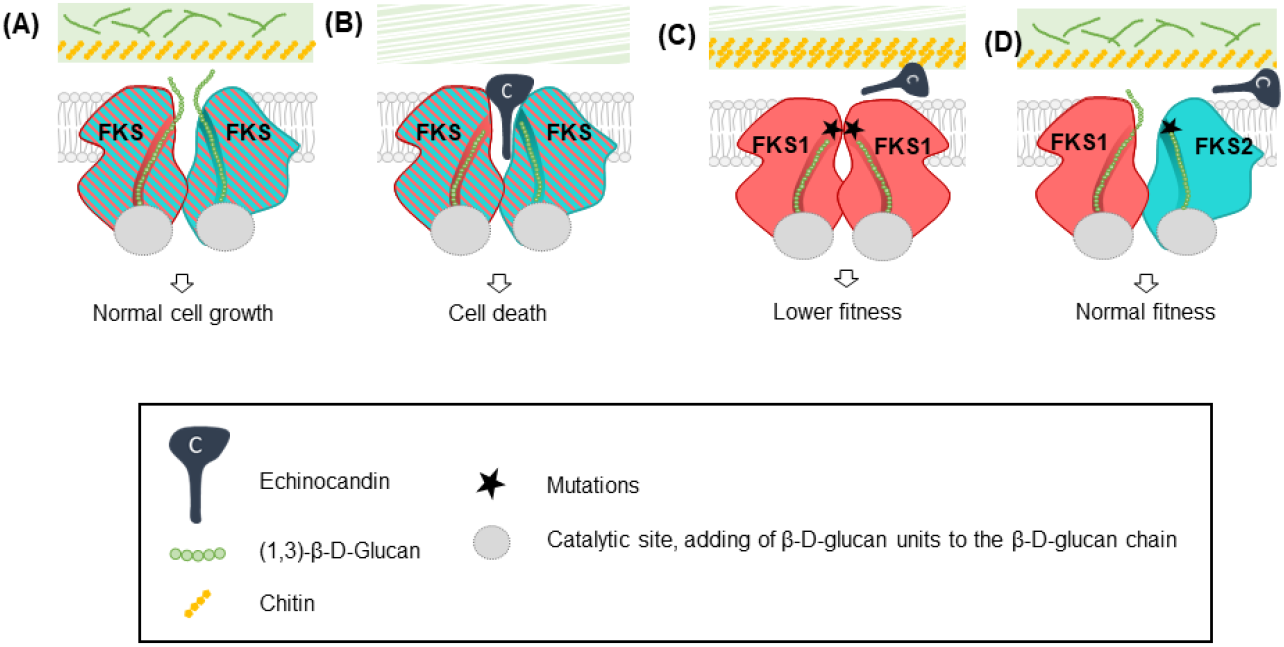
Variable fitness impact may explain the limited emergence of echinocandin resistance. (A), Wild-type strains without echinocandin treatment. (B), Wild-type strains with echinocandins. Echinocandins bind in the hydrophobic region and alter the exit of the (1,3)-β-D-Glucan chain, leading to cell death. (C), *C. albicans:* FKS1 is essential. If FKS1 is mutated, echinocandins cannot bind anymore, leading to resistance. Additionally, these mutations may alter the (1,3)-β-D-Glucan chain exit leading to a lower fitness. (D), *N. glabratus*: FKS1 and FKS2 are redundant and hypothetically forming heterocomplex. A mutation in FKS2 may alter the echinocandin binding site, thus rendering yeast resistant to this molecule. As FKS1 and FKS2 seem to be redundant in *N. glabratus*, the yeast may produce its (1,3)-β-D-Glucan chain via FKS1, thus keeping its normal fitness.

To conclude, facing an increase in echinocandins’ consumption in our center, we observed a limited change in the epidemiology of *Candida* species, characterized by a slight increase in natural-RES species, but without a significant modification for acquired-RES isolates. We speculated that this latter observation may be due to the reduced fitness of *C. albicans* strains harbouring mutations in the *fks1* gene. On the opposite, a worrying observation is the conservation of fitness in acquired RES *N. glabratus* isolates due to the functional redundancy of FKS proteins, suggesting that these isolates would be more prone to disseminate in debilitated patients. Further studies are warranted to confirm this hypothesis, notably with in-depth structural investigations using cryo-electron microscopy single particle analysis to better understand the structure-function relationship of *Saccharomycetales* FKS proteins.

## Methods

### Echinocandins consumption

The study took place in a 1500-bed University Center located in Paris (France). The consumption of echinocandin drugs (caspofungin, micafungin and anidulafungin) during the 2006-2018 period was calculated from the hospital pharmacy department database.

### Fungal epidemiology and antifungal susceptibility testing

Yeast species identification was performed thanks to the API32C auxanogram (bioMérieux) till December 2014, then relied on MALDI-TOF mass spectrometry (Bruker Daltonics). Epidemiologic trends were analysed using a multiple correlation Pearson test (GraphPad Prism8.02). Antifungal susceptibility was evaluated using the gradient diffusion method (Etest®, bioMérieux) for strains isolated from infected patients, or colonized patients with a risk of dissemination. All resistant or non-wild-type strains had their minimal inhibitory concentrations confirmed using the EUCAST broth microdilution method version e.DEF10.0 [22]. Depending on the availability of clinical breakpoints, either Clinical and Laboratory Standards Institute (CLSI) breakpoints or Etest-epidemiological cut-off [23] were used to determine patterns of susceptibility of the tested strains.

### fks sequencing and strain genotyping

*fks1* and *fks2* HS were sequenced using appropriate primers (Supplementary Table 1) [24–26]. Sequences were manually then edited and aligned using BioEdit (v.7.2.5). Clonality of isolates was confirmed using microsatellite genotyping [27–30].

### Fitness assessment

Growth curves of the isogenic susceptible and non-susceptible isolates were established in YPD media with or without a stress agent. Briefly, 10^5^ yeasts were inoculated in triplicate in 96-well plates filled with 200 μL YPD medium with or without 15 mM caffeine or 75 μg/ml Calcofluor [31]. Plates were incubated at 35°C for 48 hours in a spectrophotometer (ELX 808 BioTek) with measurement of the OD_450_. Growth curves were then drawn using the BioTek Gen5 v2.09 software.

### AlphaFold2 structural modeling

FKS1 and FKS2 models were generated with the online open-source repository ColabFold v1.5.2-patch [32] (AlphaFold2 using MMseqs2) after entering the corresponding protein sequences. Graphical representations were done with PyMol Molecular Graphics System, Version 1.3r1 (Schrödinger, LLC) and ChimeraX [33].

### Ethics approval

Specimens were collected through routine clinical tests and patient-identifiable information was anonymized. As recommended in the ethical standards of the French Ethics Committee on human experimentation and with the Helsinki Declaration of 1975, as revised in 2008, no written or verbal informed consent to participate in this study from the patient was necessary.

## Author contributions

J.G. and C.H. conceptualized and designed experiments; J.G. supervised the studies; V.L. and E.D. performed the Eucast testing, M.A. evaluated the echinocandin consumption, J.B. retrieved the fungal epidemiology, S.V. performed Etest and *Candida* identification, Y.K. performed *fks* and strain genotyping as well as fitness analysis, A.G. and S.P. performed structural modelling and analysis of FKS mutations; J.G. and A.G. wrote the manuscript with input from all authors. All authors approved the draft before submission.

## Acknowledgements

The authors want to thank Manon Ruffin for her help in graphical art and Pierre Vincent for technical help.

## Appendix. Supplementary materials

